# A retrospective cluster analysis of COVID-19 cases by county

**DOI:** 10.1101/2020.11.12.379537

**Authors:** Fadel M. Megahed, L. Allison Jones-Farmer, Steven E. Rigdon

## Abstract

The COVID-19 pandemic in the U.S. has exhibited distinct waves, the first beginning in March 2020, the second beginning in early June, and additional waves currently emerging. Paradoxically, almost no county has exhibited this multi-wave pattern. We aim to answer three research questions: (1) How many distinct clusters of counties exhibit similar COVID-19 patterns in the time-series of daily confirmed cases?; (2) What is the geographic distribution of the counties within each cluster? and (3) Are county-level demographic, socioeconomic and political variables associated with the COVID-19 case patterns? We analyzed data from counties in the U.S. from March 1 to October 24, 2020. Time series clustering identified clusters in the daily confirmed cases of COVID-19. An explanatory model was used to identify demographic, socioeconomic and political variables associated the cluster patterns. Four patterns were identified from the timing of the outbreaks including counties experiencing a spring, an early summer, a late summer, and a fall outbreak. Several county-level demographic, socioeconomic, and political variables showed significant associations with the identified clusters. The timing of the outbreak is related both to the geographic location within the U.S. and several variables including age, poverty distribution, and political association. These results show that the reported pattern of cases in the U.S. is observed through aggregation of the COVID-19 cases, suggesting that local trends may be more informative. The timing of the outbreak varies by county, and is associated with important demographic, socioeconomic and geographic factors.

## Introduction

When considering the aggregated number of cases in the U.S. as a whole, the COVID-19 pandemic has so far exhibited distinct waves. The first wave began in March 2020, peaked in April, and then receded somewhat following a widespread lockdown. The number of cases began to rise again in early June once states began to reopen. With further restrictions and health guidelines, the cases seemed to recede by the end of July; however, additional cases are currently emerging. In terms of number of reported COVID-19 cases, the second wave was larger than the first. Paradoxically, almost no county in the U.S. has exhibited the multi-wave pattern seen in the aggregated U.S. data. Many counties, especially in the Northeast, exhibited a large first wave followed by a smaller second wave. On the other hand, many counties in the Midwest exhibited a small first wave followed by a larger second wave in terms of cases. Some counties, especially small rural ones, have not yet exhibited clear peaks in the number of reported COVID-19 cases (as of this writing).

Figure 1 shows the 7-day moving averages for the number of new cases in the U.S. and eight individual counties. For example, in New York, NY, we see a large early wave. In Madison County, IL (a Midwestern county near St. Louis, MO), a large sustained increase in cases is observed in the late summer. In Butler County, Ohio there is an increasing trend in confirmed cases beginning in late summer, while in Falls Church, Virginia, there is only a small number of confirmed cases with no observable pattern.

**Figure 1.**
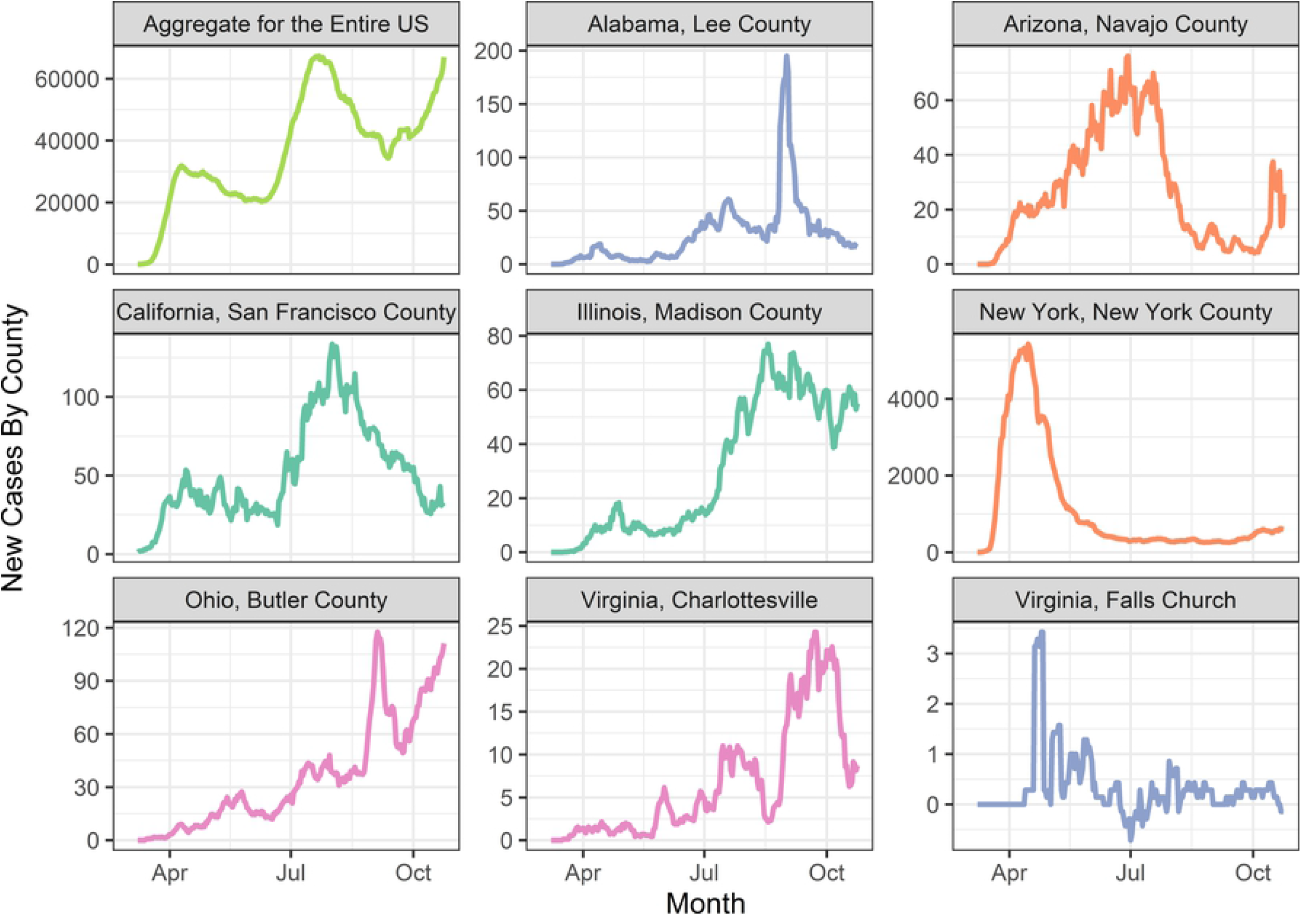
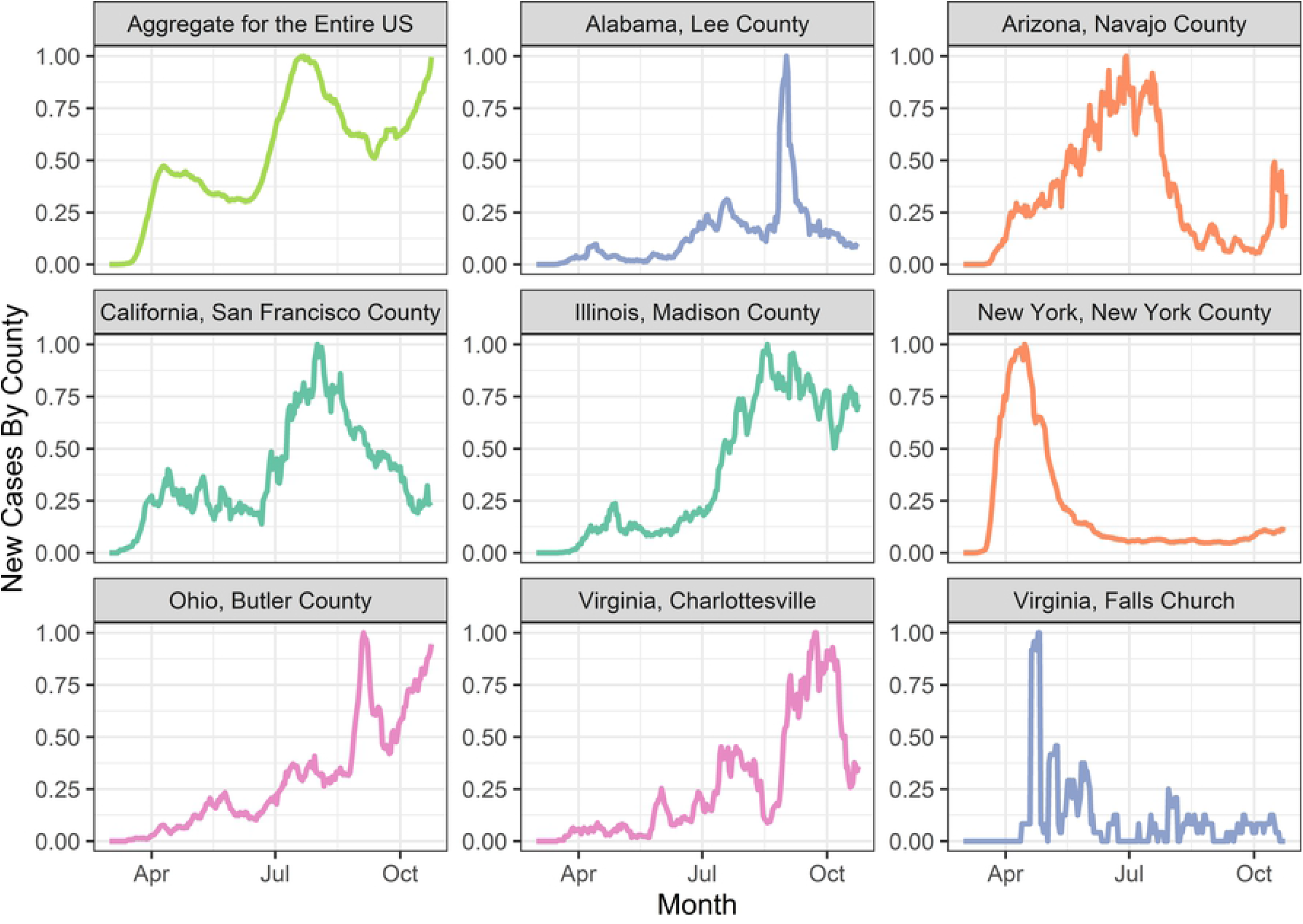
The time series profiles for the entire U.S. and the sample of eight counties. The number of cases varied greatly among these counties. To assess the shape, we scaled each time series so that the maximum value is 1.

The observed local-level patterns can be used to inform decisions made to mitigate the pandemic. The roles of the federal and local governments in enacting measures like school closures and business restrictions to combat the virus have been under debate. Not everyone agrees on how decisions should be made. For example, Koh^1^ argues that there should be one strategy, not 50. Similarly, Haffajee and Mello [2] argue that “strong, decisive national action is therefore imperative.” On the other side of the debate, others, such as Davidson [3] argue that the federal government does not have the authority enact measures like lockdowns, because these powers are reserved for the states. The arguments for or against federal vs. state vs. local control of mitigation standards may be clarified by a better understanding of the pattern of outbreaks.

## OBJECTIVE

The observation that almost no counties follow the pattern of the aggregated number of reported COVID-19 cases in the U.S. led us to pose these research questions:

1. How many distinct clusters of counties exhibit similar COVID-19 patterns in the time-series of daily confirmed cases?
2. What is the geographic distribution of the counties within each cluster?
3. Are county-level demographic, socioeconomic and political variables associated with the COVID-19 case patterns?

To explore these research questions we used a time series cluster analysis of counties within the contiguous U.S. to identify groups of counties with similar COVID-19 outbreak patterns. A visualization of the cluster solution provides information on the distribution of the cluster patterns across the U.S. Finally, we used multinomial logistic regression to develop an explanatory model that identifies county-level variables that are associated with some of the observable variation in the COVID-19 outbreak patterns.

## MATERIALS AND METHODS

This analysis was conducted in three stages. In Stage 0, county-level data were gathered from several sources, merged, and preprocessed for consistency. In Stage 1, time series clustering was performed on the number of newly reported confirmed COVID 19 cases per day. Finally, in Stage 2, an explanatory model was fit to describe the relationship between cluster membership and several socioeconomic and political factors describing the counties. An overview of the methods is given in Figure 2.

**Figure 2.**
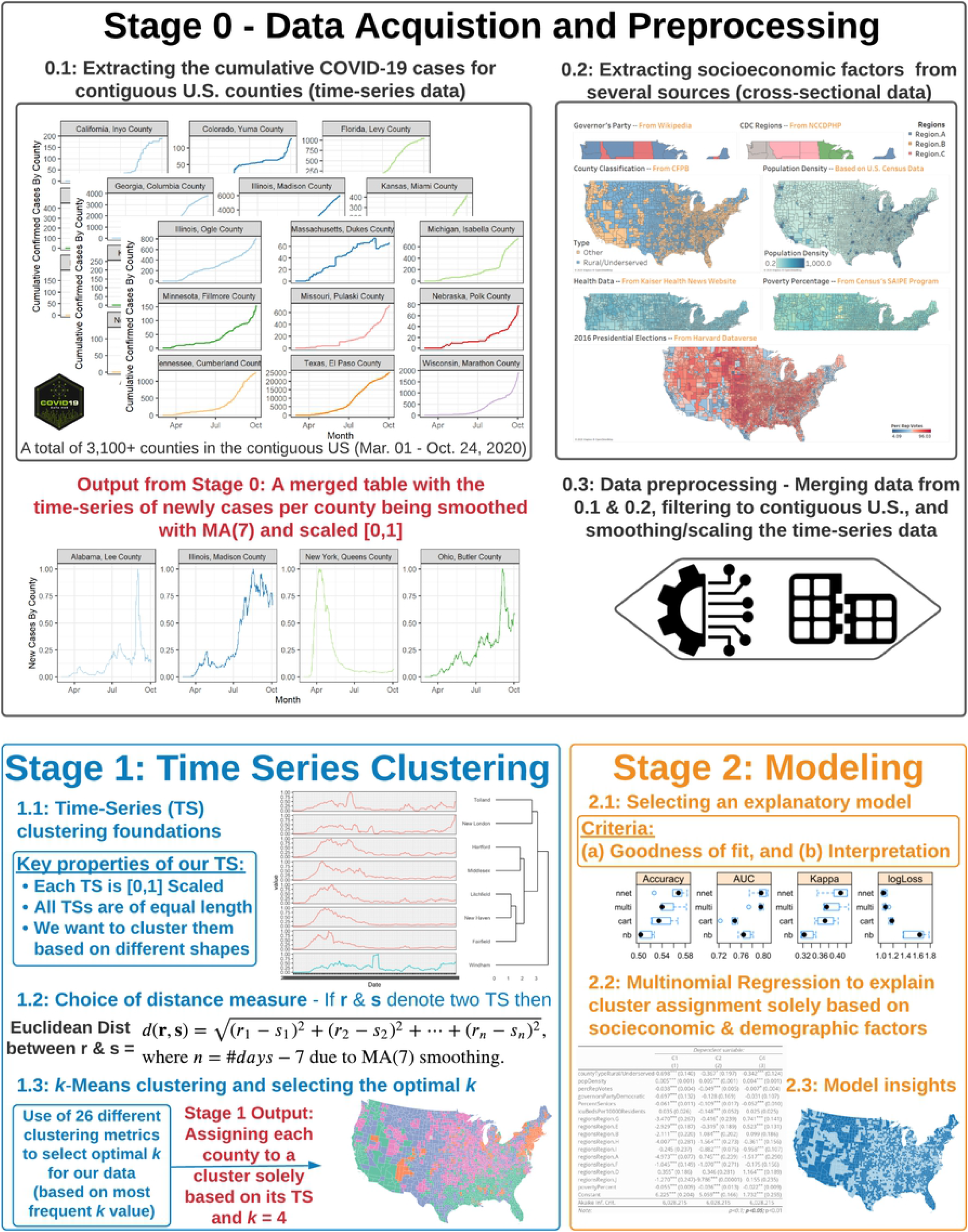
An overview of the three methodological stages for this study: data acquisition and preprocessing, time series clustering; and modeling

### Stage 0: Data Acquisition and Preprocessing

Guidoti and Ardia [4] provide an open-source COVID-19 data hub to facilitate research regarding the novel coronavirus. The disease outbreak was declared a pandemic by the World Health Organization (WHO) on March 11, 2020 and a national emergency by the U.S. on March 13, 2020. To capture the progression of disease in the U.S., the number of confirmed COVID-19 cases at the county level from March 1, 2020 through October 24, 2020 was extracted from the COVID-19 data hub [4]. Only data from the contiguous 48 states were included. The number of newly reported confirmed COVID-19 cases per day is used to establish the pattern of the epidemic progression in each county, and is the only information used to determine the time series clustering of the counties.

Additional county-level exogenous variables were extracted from several sources to be used in an explanatory model to describe the clusters of outbreak patterns. The variables include the following.

- Rural/Underserved Counties: The Consumer Financial Bureau^6^ provides a list of rural or underserved counties based on the Bureau’s interpretive rule (Regulation Z) using data from the Home Mortgage Disclosure Act [5].
- Population Density: Based on the US Census Data [6], the land area in square miles for each county was extracted. This was combined with the population in each county to compute the county’s population density.
- Voting Results: The MIT Election and Data Science Lab [7] provides voting results for all counties in the 2016 Presidential election. This data was used to compute the percentage of total votes for President Trump.
- Governor’s Party Affiliation: Based on Wikipedia’s Table of State Governors [8], the Governor’s political party affiliation was obtained. Given that the District of Columbia does not have a governor, a value of “Democratic” was imputed since D.C.’s Mayor is a Democrat.
- Health Variables: From Kaiser Health News [9], county-level information was extracted on: (a) number of ICU beds per 10,000 residents; (b) percent of population aged 60+.
- Region: Using the Center for Disease Control and Prevention’s (CDC) 10 Region Framework for Chronic Disease Prevention and Health Promotion [10], geographic region indicators were obtained. Figure 3 shows these ten regions. This source of defining regions within the U.S. was selected because the “CDC’s National Center for Chronic Disease Prevention and Health Promotion (NCCDPHP) is strengthening the consistency and quality of the guidance, communications, and technical assistance provided to states to improve coordination across our state programs” [10].
- Poverty: Based on the U.S. Census’s Small Area Income and Poverty Estimates (SAIPE) Program [11], the percent of population in poverty for each county was extracted. The estimate is based on 2018 data (released in December 2019). At the time of the start of our analysis, these estimates were the most up to date publicly available data.

**Figure 3.**
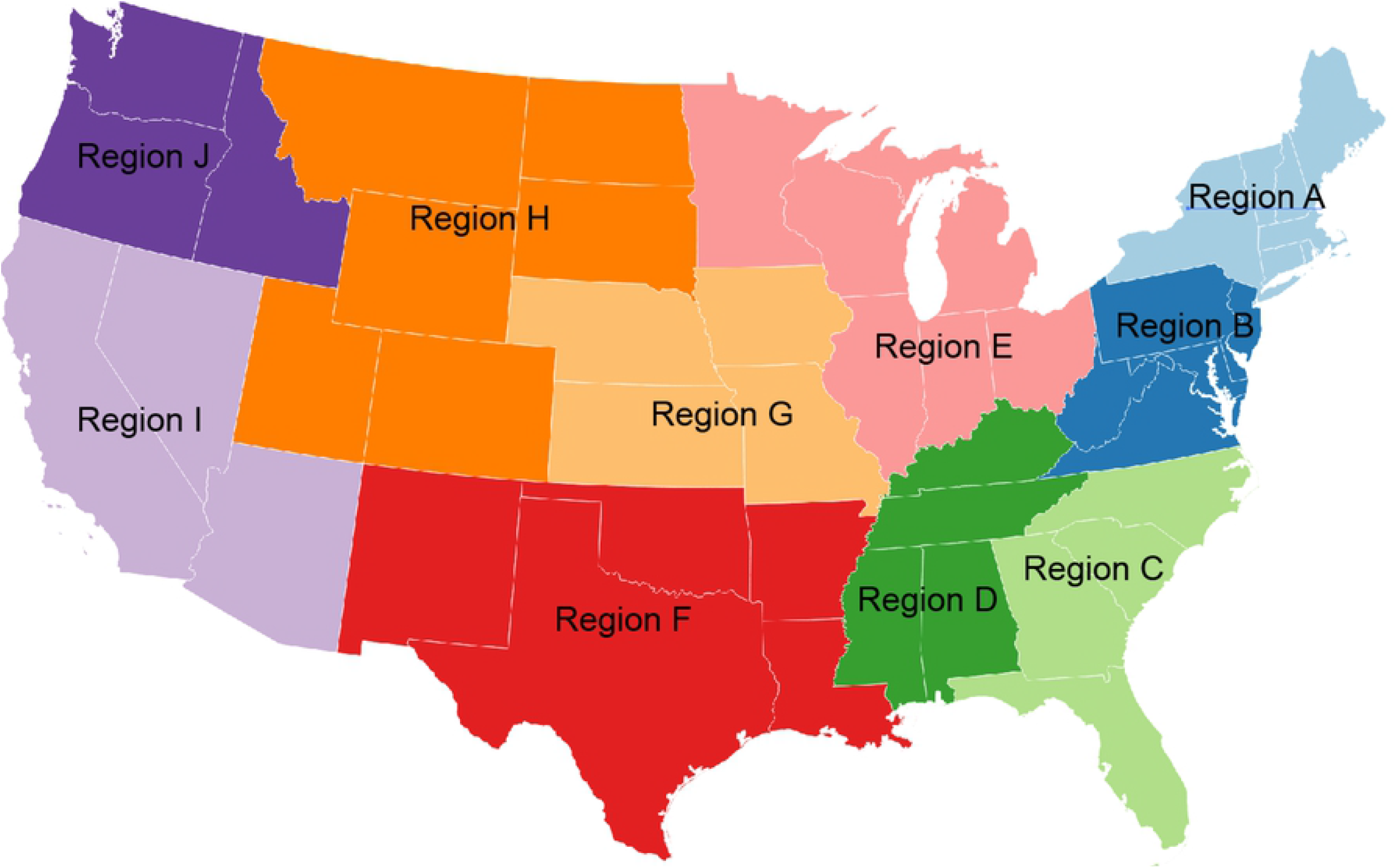
The CDC’s ten regions

### Stage 1: Time Series Clustering

The cluster solution is based on the daily number of newly reported, confirmed COVID-19 cases by county over time. No other information is used to determine the cluster membership. For time series clustering, there are three important decisions that affect the cluster solutions: (1) the scaling and preprocessing of the data; (2) the distance measure between clusters; and (3) the clustering algorithm. Liao^13^ provides an accessible overview of time series clustering methods.

#### Scaling the data

For each county, the counts of newly reported confirmed COVID-19 cases were smoothed using a seven-day moving average to account for the weekly seasonal pattern. We rescaled the seven-day moving averages for each county such that the values fall between 0 and 1 as follows.

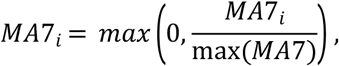

where *MA7* is the vector of all seven-day moving averages of newly reported confirmed COVID-19 cases for a given county and *MA7_i_* is an observation from this vector. Some counties reported negative cases on some days, resulting from reclassification of their recent cases: thus, it is possible that the moving average is negative in a few cases. This is why the max is required in the above equation. Rescaling the data in this way is done to focus the time series clustering on the pattern of the disease progression, not the magnitude of the daily case counts which may be dependent many factors such as county size, population, etc.

#### Distance Measure

In order to cluster the scaled time series profiles, it is necessary to measure the distance between the profiles. There are many ways to measure the distance between time series, including Euclidean distance, dynamic time warping [13], Pearson’s correlation coefficient and others [12,14]. For this analysis, Euclidean distance was chosen specifically because it is computationally efficient and is not an elastic measure such as dynamic time warping. Euclidean distance provides a shape-based measure that gives one-to-one indexing across the profiles; thus, it preserves the time-based shape of the profiles which is desired in this application of time series clustering. The Euclidean distance between two time series, *r* and *s*, of length *T* is defined as

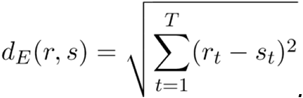

#### Clustering Algorithm

A large number of clustering algorithms have been proposed in the literature, which have been studied and compared in the context of time series [12,14]. For this analysis, *k*-means clustering was used. A possible limitation of *k*-means clustering approach for exploratory research is the number of clusters must be pre-determined. In addition, there have been many measures proposed to assess the validity of the cluster solution [15]. The R package, NbClust [16] computes up to 30 cluster validity indices for cluster solutions of a variety of sizes. This allows the analyst to select the solution with the most homogeneity within the clusters, and provides a systematic method for selecting the optimal number of clusters in a data set. For this analysis the NbClust package was used with the *k*-means clustering method to select the optimal number of clusters.

### Stage 2: Modeling

The clustering method described above resulted in each county being assigned to one of four different clusters. An explanatory/predictive model that includes the county-level exogenous variables was fit both to describe the relationship between these variables and to predict cluster membership. Specifically, a multinomial regression analysis was fit because it balanced computational efficiency, predictive accuracy and ease of explanation. Prior to the selection of the multinomial regression analysis, the use of a classification and regression tree (CART), a naive Bayes (NB) classification method, and a model averaged neural network (avNN) representing a typical ensemble structure was considered. Five-fold cross-validation with bootstrap sampling was used for model selection. The results of the model comparison from a 20% holdout sample showed nearly equivalent predictive performance between the avNN and the multinomial regression with average model accuracy of 0.677 and 0.666, respectively.

## Results

### Time Series Cluster Solution

We computed *k*-means cluster solutions ranging from *k* = 2 to 50 clusters based on the scaled time series profiles of the daily confirmed COVID-19 cases. We evaluated twenty-six recommended cluster validity indices [15] for each *k*. For a full list of the cluster validity indices considered, see Charrad et al. [16]. Eight of the twenty-six cluster validity indices indicate a four-cluster solution is preferred. The second most preferred solution is a two-cluster solution, which was selected by seven out of the twenty-six indices. Based on majority rule of the validity indices, a four-cluster solution is retained. The geographic distribution of the clusters is shown in Figure 4. Figure 5 shows the time series profiles for each of the clusters.

**Figure 4.**
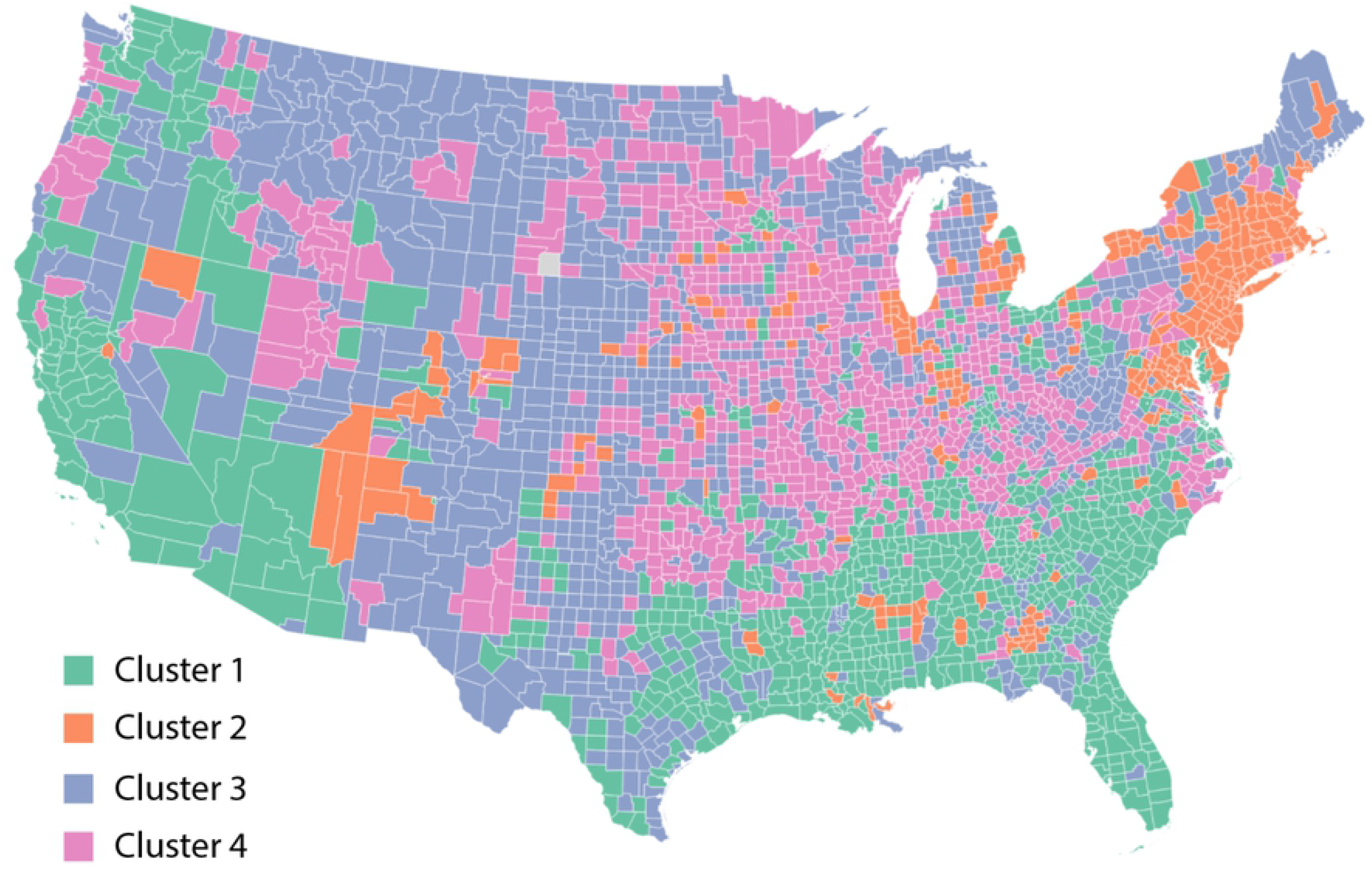
Map of the four scaled time-series profile clusters by county in the contiguous United States

**Figure 5.**
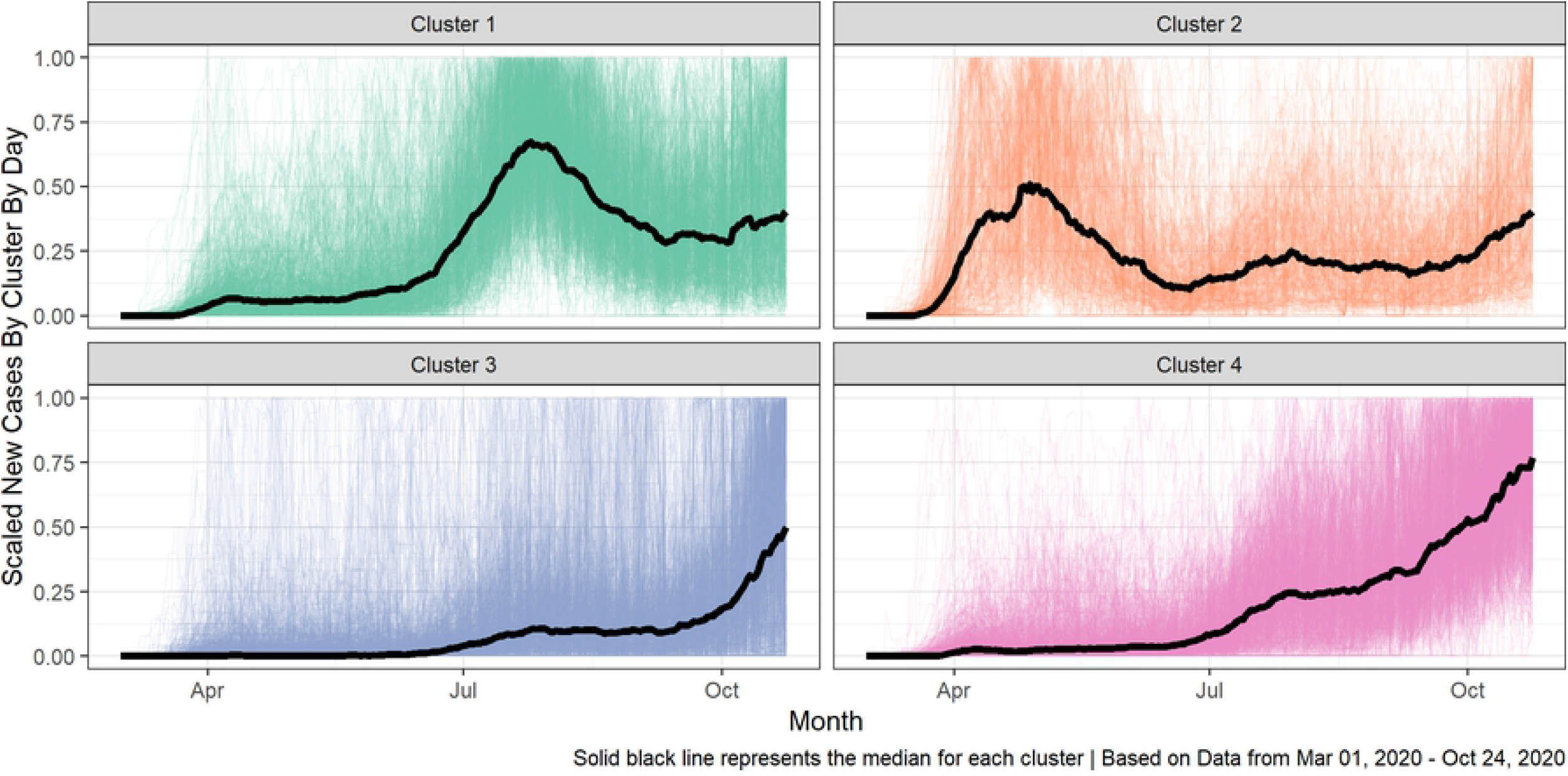
A spaghetti plot, where the median scaled time-series profile for each cluster is bolded and the remaining profiles within the cluster are shown in the background

### Modeling Results

In this section, we provide the results of an explanatory model to answer our third research question: Are county-level demographic, socioeconomic and political variables associated with the COVID-19 case patterns? We also use this model in a predictive manner to support the validity of our cluster solution.

### Model Explanation

The coefficients from the multinomial logistic regression are given in Table 1. The baseline category for analysis was chosen to be the cluster with the largest number of counties, which is Cluster 3 (C3). Each coefficient shows the linear change in the natural log of the odds ratio of a county classifying in the corresponding cluster as indicated by the column vs. the baseline cluster.

**Table 1:** Results of multinomial logistic regression for the probability of falling in C1, C2, and C4. Note that we have used C3 as the reference cluster since it contained the largest number of counties.

|  | Cluster |  |  |  |  |  |
| --- | --- | --- | --- | --- | --- | --- |
|  | C1 |  | C2 |  | C4 |  |
| | $\beta$ | $\exp(\beta)$ | $\beta$ | $\exp(\beta)$ | $\beta$ | $\exp(\beta)$ |
| County: Rural/Underserved | -0.698*** | 0.498 | -0.367* | 0.693 | -0.342*** | 0.710 |
| Population Density | 0.005*** | 1.005 | 0.005*** | 1.005 | 0.004*** | 1.004 |
| Percent Republican Votes | -0.038*** | 0.963 | -0.049*** | 0.952 | -0.007* | 0.993 |
| Governor Party Affiliation | -0.697*** | 0.498 | -0.128 | 0.88 | -0.031 | 0.969 |
| Percent Seniors | -0.061*** | 0.941 | -0.109*** | 0.897 | -0.052*** | 0.949 |
| ICU Beds/10,000 residents | 0.035 | 1.036 | -0.148*** | 0.862 | 0.025 | 1.025 |
| Region A | -4.973*** | 0.007 | 0.745*** | 2.106 | -1.517*** | 0.219 |
| Region B | -2.111*** | 0.121 | 1.084*** | 2.956 | 0.099 | 1.104 |
| Region D | 0.355* | 1.426 | 0.346 | 1.413 | 1.164*** | 3.203 |
| Region E | -2.929*** | 0.053 | -0.319* | 0.727 | 0.523*** | 1.687 |
| Region F | -1.045*** | 0.352 | -1.070*** | 0.343 | -0.175 | 0.839 |
| Region G | -3.470*** | 0.031 | -0.416* | 0.660 | 0.741*** | 2.098 |
| Region H | -4.007*** | 0.018 | -1.564*** | 0.209 | -0.361** | 0.697 |
| Region I | -0.245 | 0.783 | -0.882*** | 0.414 | -0.958*** | 0.384 |
| Region J | -1.270*** | 0.281 | -9.786*** | 0.000 | 0.155 | 1.168 |
| Poverty Percent | -0.055*** | 0.946 | -0.036*** | 0.965 | -0.022** | 0.978 |
| Constant | 6.225*** | 505.223 | 5.059*** | 157.433 | 1.732*** | 5.652 |

Notes: The baseline cluster is C3; \* $p < 0.1$ , \*\* $p < 0.05$ , \*\*\* $p < 0.01$

For example, in the column of Table 1, labeled C1, the first coefficient corresponds to the variable “County Type Rural/Underserved.” The coefficient, −0.698 (*p* < 0.01), suggests that, *ceteris paribus*, the odds, or relative risk of a county classifying in C1 vs. C3 is lower by a factor of exp(-.725) = 0.498 when that county is Rural/Underserved.

### Cluster Validity

Our *k*-means solution meets the five recommended guidelines for reporting cluster analysis solutions given by Clatworthy et al. [17], who recommend reporting the computer program, similarity measure, clustering algorithm, decision criterion for number of clusters, and an evaluation of cluster validity. Our methods and results have provided the first four of these. In addition, we provide evidence of cluster validity by developing an explanatory/predictive model exploring the relationship between exogenous variables and cluster membership. Table 2 shows the counts and percent accuracy of predictions for each of the clusters based on the explanatory/predictive model. For each cluster, the largest predicted category is the true cluster; however, some clusters were easier to identify than others. Figure 6 shows the geographic distribution of the classification accuracy based on the explanatory/predictive model. Counties correctly classified in from the model are shown in dark blue, while those incorrectly classified are shown in light blue.

**Table 2:**
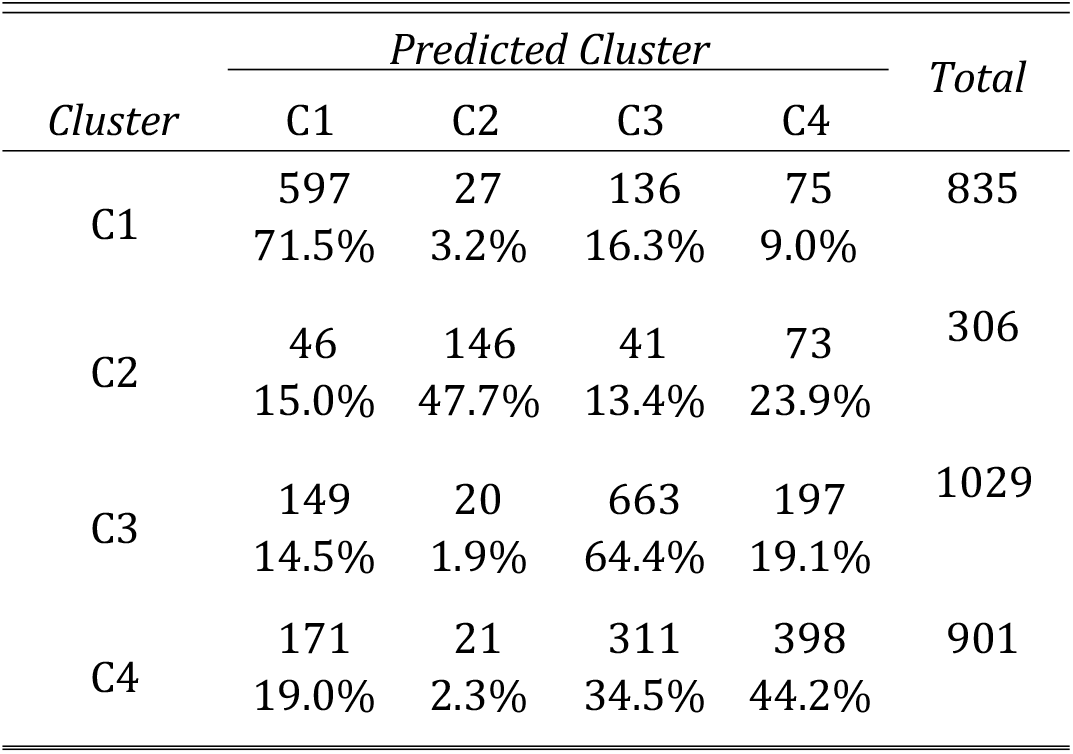
A summary of the predictive performance of the multinomial regression model. For a given cluster, the first and second rows show the number and percentage of predicted cases, respectively.

**Figure 6.**
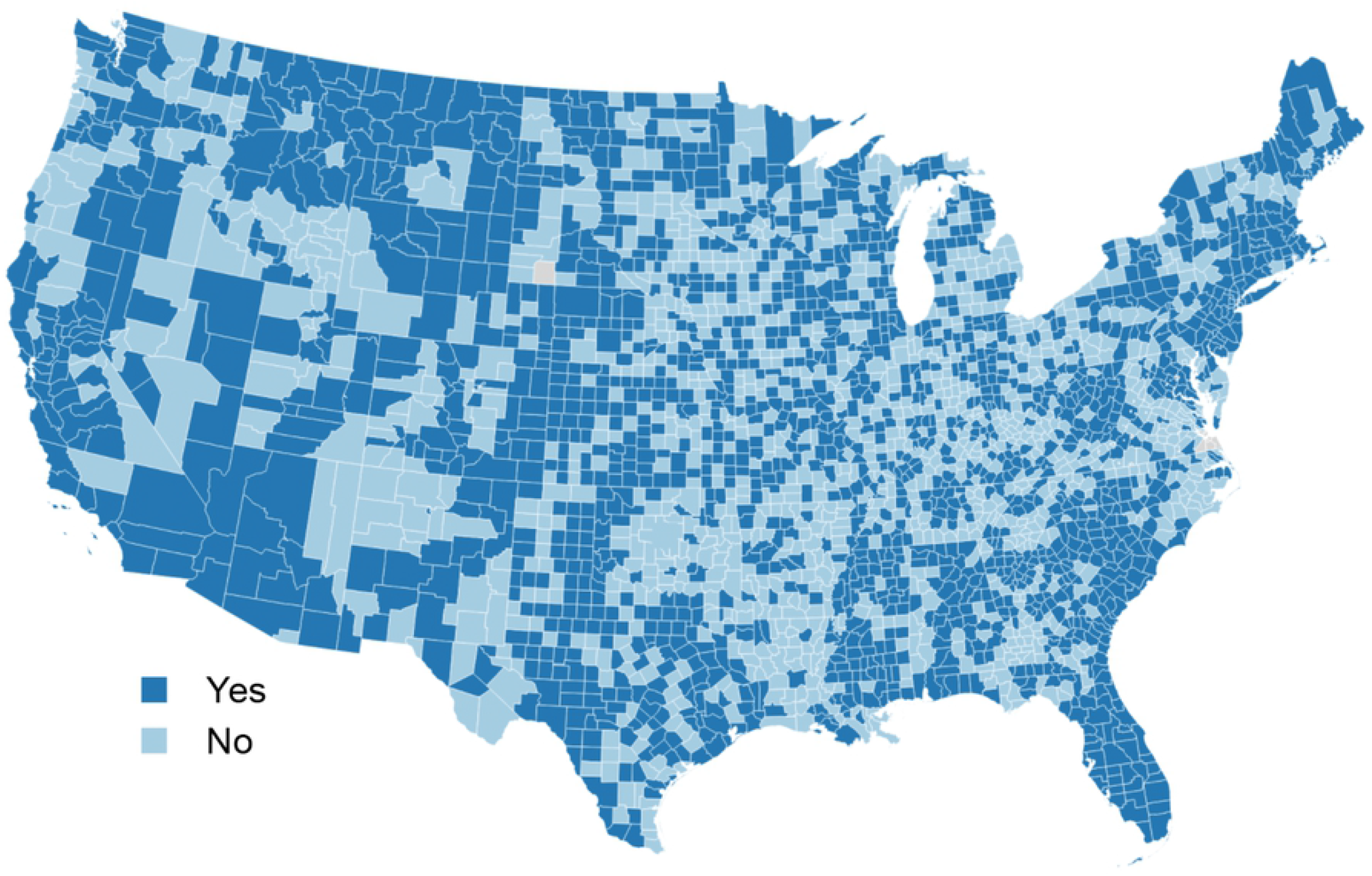
Map of the prediction accuracy of the multinomial logistic model describing the time series cluster solution

## Discussion

The results of the cluster analysis provide an answer to our first research question: How many distinct clusters of counties exhibit similar COVID-19 patterns in the time-series of daily confirmed cases. Based on the time series cluster analysis, our results suggest that there are four distinct cluster patterns.

Figure 4 provides a map of the *k*-means four-cluster solution of the scaled time series profiles across the U.S. This map gives further insight into the geographic distribution of the clusters, which addresses our second research question: What is the geographic distribution of the counties within each cluster?

The geographic distribution shows that C1 is primarily centralized in the Southern and Western U.S. states. Figure 5 gives a visual summary of the cluster solutions. From this plot we see that most counties clustering in C1 experienced an outbreak that began in late summer. Counties in C1 include San Francisco County, California and Madison County, Illinois.

The counties clustering in C2 are located distinctly in the upper East Coast with a few counties in the western U.S., the South and Midwest. From Figure 5 it is clear that these counties experienced an early outbreak of COVID during the spring. Counties in C1 include, e.g., New York County, NY and Navajo County, AZ.

The largest number of counties clustered in C3. These are located throughout the western states and in rural areas, sporadically throughout the U.S. Counties in C3 include Lee County, Alabama and Falls Church, Virginia. The pattern of the outbreak in these counties seems to be increasing beginning in the late fall.

Finally, the counties that clustered with C4 are located primarily throughout the Midwest, but include a few counties on the East Coast and in the western U.S. Counties in C4 include, e.g., Butler County, Ohio and Charlottesville, Virginia. These counties experienced a late summer outbreak that seems to have been sustained.

The third research question is: Are county-level demographic, socioeconomic and political variables associated with the COVID-19 case patterns? The results of the multinomial regression model suggest that county-level geographic, political, and socioeconomic factors are significantly associated with the four observed COVID-19 case patterns. Comparing the C1 (early summer outbreak) to the C3 (the baseline, late fall outbreak), counties with higher population density are associated with a slightly higher relative risk of classifying in C1 (*β* =.005, *p* < 0.01). Location in a rural county, a higher percentage of Republican votes in the 2016 presidential election, a Democratic governor, a higher percentage of Seniors, location in any CDC Region other than I (California, Nevada, Arizona) and D (Central southern states), and a higher poverty percentage are associated with a lower relative risk of a county classifying in C1.

Comparing the C2 (early spring outbreak) to C3 (the baseline, late fall outbreak), increased population density, and location in CDC region A or B (both in the Northeast) are associated with an increased relative risk of classifying in C2. Percent of Republican votes in the 2016 presidential election, percent of seniors, the number of ICU beds, location in regions F,H, I, or J (all western regions), and higher poverty percentage are all associated with a lower relative risk of classifying in C2. Recall that region C2 consists of counties such as New York that experienced the early spring outbreak of COVID. Many counties that experienced large early outbreaks are high density urban counties.

The differences between C4 and C3 are more subtle. At present, the timing of the outbreak in these clusters differs only by a matter of weeks, and the full pattern is emerging. There are, however, some observed differences in some of the demographic, political, and socioeconomic variables between these clusters. For example, the relative risk of classifying in C4 is lower for rural counties (*β* = −.342, *p* < 0.01), and there is a slightly lower risk associated with a higher percentage of seniors (*β* = −0.052, *p* < 0.01) and a higher poverty percentage (*β* = −0.022, *p* < 0.05. Location in CDC regions A (Northeast), H (Upper Western States) and I (California, Nevada, Arizona) also lowers the relative risk of classifying in C4 (the late summer outbreak) vs. C3 (late fall outbreak). Specifically, location in Regions D, E, and G (regions in the central South and Midwest) along with a higher population density are related to increased relative risk of classification in C4 vs. C3.

Figure 6 shows the geographic distribution of the accuracy of the multinomial logistic model in predicting cluster membership. Counties that are accurately predicted are shown in dark blue, while counties that are not accurately predicted by the model are shown in light blue. From this map it is clear that additional data are needed to describe the outbreak patterns throughout the U.S.

## Conclusion

The local patterns in outbreaks suggest that decisions regarding the timing of mitigation efforts should be informed by local conditions. Local conditions vary across the country and even within each state, and the clusters of patterns exhibit spatial distributions. By understanding the patterns of COVID-19 progression across the country, policy and mitigation standards can benefit from regional information at a given time in order to better preserve public health.

## Acknowledgements

None

## SUPPORTING INFORMATION

The R statistical software system, version 4.0.2 was used for all processing and analysis of data. To facilitate the reproduction of our research and to follow the best practices of Jalali *et al*. ^18^ in reporting and documenting analyses for COVID-19, we have capitalized on *R Markdown* documentation mechanism to produce an automated report that contains all our data, analysis and results, which we host at https://fmegahed.github.io/covidanalysis.html.

